# Exploration of diuretic potential of different parts (flower, bark, leaf) of *Butea monosperma*

**DOI:** 10.64898/2025.12.22.696018

**Authors:** Nafisa Lubna, Sonia Akter, Bidduth Kumar Sarker, Md. Asif Mahmud, Md Masuk Ur Rashid, Sadia Afrin, Sad Al Rezwan Soumik, Mala Khan, Md. Ibrahim Khalil

**Author notes:** Correspondance: Md. Ibrahim Khalil,; Mala Khan.

## Abstract

Hypertension (HTN) or high blood pressure poses major global health concerns, and the conventional synthetic antihypertensive drugs are effective, though their long-term use is often limited due to adverse effects. *Butea monosperma* (*B. monosperma*), commonly known as the Flame of the Forest, the traditional use as medicinal plant for diuresis and other usage in rural areas in South Asia suggests a role for its antihypertensive potential. The study aimed, first to identify the diuretic constituents present from different parts (flower, bark, and leaf) of *B. monosperma* and to evaluate their diuretic activity through *in vivo* experimentation. The chemical profiling of the extracts of different parts were performed using Liquid Chromatography Mass Spectrometry (LC–MS/MS), while diuretic efficacy was assessed in Swiss Albino wistar rats. Urinary volume and electrolyte excretion (Na□, K□, and Cl□) were measured to elucidate the possible mechanism of action. Statistical analyses were conducted using one-way ANOVA followed by multiple comparison test, with significance set at *p* < 0.05. LC MS/MS analysis confirmed the presence of chlorthalidone, a thiazide-like diuretic compound, in all three plant parts under selective ionization mode. In vivo results demonstrated a significant, dose dependent increase in urine output and electrolyte excretion in treated rats compared to controls (*p* < 0.05). These findings provide both phytochemical and pharmacological evidence that *B. monosperma* possesses potent diuretic activity, supporting its traditional use and highlighting its potential as a natural source for the development of novel antihypertensive and diuretic therapeutics.

## Introduction

The variety of molecules found in the natural flora and fauna, particularly in plants from different locations, have been proven to contain medicinal properties and the potential to combat complex diseases. Scientists have long focused on the isolation of bioactive compounds from plants as plants are rich in a wide range of secondary metabolites such as flavonoids, alkaloids, tannins, and terpenoids (Twaij & Hasan, 2022). *Butea monosperma (B. monosperma),* commonly known as the “Flame of the Forest,” belongs to the Fabaceae family. It is also referred to by various local names including Astesu, Palash, Mutthuga, Bijasneha, Dhak, Khakara, Chichra, Bastard teak, and Bengal kino, as reported by local people and tribes across the Indian subcontinent(Kirtikar et al., 1935).

Hypertension (HTN), often known as high blood pressure, is a medical condition that causes significant mortality worldwide And is has also been associated with a higher relative risk of both cardiovascular and renal disease including coronary heart disease, stroke, congestive heart failure, and renal insufficiency (He & Whelton, 1999). A 5-6 mm Hg lower level of diastolic pressure is typically associated with 20-25% lower risk of incident coronary heart disease (Stamler J, 1993, Collins R, 1990, Whelton PK, 1994).

However, its initiation is complex, not caused by a single issue but by a network of factors, including obesity, alcohol intake, smoking tobacco, salt-heavy diet and lack of exercise, all has a progressive linked to hypertension (Osoro & Rajanandh, 2025). These factors induce overactivity in the body’s core regulatory systems, specifically the sympathetic nervous system and the renin-angiotensin-aldosterone system (RAAS), which drive blood pressure up by constricting blood vessels and retaining salt and water (Esler, 2015). Treatment for hypertension is important as it turns on other irreversible outcomes, for example, cardiovascular disease, and the progressive functional degradation of the kidneys, leading to chronic kidney disease (Lackland, Voeks, & Boan, 2016). Changes in lifestyle, for example eating healthy diet, daily physical exercise, efforts for loosening weight etc, are the most effective preventative and foundational way to keep blood pressure in check (Whelton et al., 2018). However, for many patients, medications are needed to avoid risk outcomes. With medication, in parallel with lifestyle efforts, a protective measure encores the long-term benefits of diet and exercise take effect (Whelton et al., 2018).

As a matter of fact, numbers of synthetic antihypertensive medications are available to treat hypertension but in terms of efficacy, variations are noticeable and might cause side effects like dry mouth, dizziness, mental discomfort, gastrointestinal disturbances, and vision abnormalities, among other things (Gebreyohannes EA, Bhagavathula AS, Abebe TB, et al., 2019). These unfavourable side effects have a significant impact on overall health and quality of life. Diuretics are defined as drugs increasing the excretion of water and electrolytes from the patients’ bloodstream through the kidneys and thereby elevating the rate of urine flow for immediate relief (add citation). According to standard textbooks in the field of pharmacology, diuretics are used in the treatment of edema, heart failure, or hypertension (Aktories et al. 2017; Buckingham 2020). The compounds exhibiting diuretic effects are of five classes : Carbonic anhydrase inhibitors (CAIs), Loop diuretics, Osmotic diuretics, Potassium-sparing diuretics, and Thiazides (Lüllmann et al. 2016). Thiazides are compounds, make the kidneys pull salt and extra water into your pee. i.e: Chlorthiazide, Hydrochlorothiazide, chlorthalidone etc and loop diuretics (e.g furosemide or bumetanide) affect part of the kidneys (the loop of Henle) to get salt and excess water out of your body (Sica, Carter, Cushman, & Hamm, 2011). On the other hand, potassium-sparing diuretics such as triamterene or amiloride don’t let salt and water lose too much potassium in the process. Diuretics, especially thiazides were the first tolerated efficient antihypertensive drugs that significantly reduced cardiovascular morbidity and mortality in placebo-controlled clinical studies but recent meta-analysis of placebo-controlled trials suggested that the beneficial effects of thiazide diuretics could be a class effect (Salvetti A and Ghiadoni L, 2006).

Over the past three decades, significant efforts have been made to explore alternative plant-based medications with hypotensive and antihypertensive potential. It should be noted that the prevalence of hypertension, as well as the availability and efficacy of medications, depends on factors such as age, race, gender, living conditions, and socioeconomic status; especially for chronic health conditions like hypertension. Despite the availability of synthetic drugs, a large proportion of the population in low-income countries continues to rely on traditional medicinal practices. As hypertension is a global health problem, a lot of this population would be interested in having an effective natural remedy for hypertension that has less health impacts than other prescribed synthetic treatments.

Medicinal plants have been widely recognized for their potential to combat hypertension and serve as a natural alternative to synthetic drugs, a consideration that also applies to *B. monosperma* (Sultana S, Asif HM, 2017). To date, no scientific investigation has been conducted on the therapeutic potential of *B. monosperma* aqueous methanolic extract as a diuretic to battle hypertension immediate blood pressure control. As a medicinal plant, *B. monosperma* could contain numerous bioactive compounds that may contribute to its antihypertensive effects; the diuretic potential remains poorly investigated and evaluated. While many plants show in vivo diuretic activity, including increased urine volume and renal electrolyte excretion, the mechanisms, dose-response, and efficacy of *B. monosperma* have not been fully validated in modern animal models. This study aims to identify any medicinally established diuretic compounds from different extracts of *B. monosperma* and evaluate their diuretic functional activity with dose dependent procedure, bridging traditional knowledge with scientific validation.

## 2. Materials and Methods

### 2.1. General description of *B. monosperma*

*B. monosperma* is a medium-sized deciduous tree of the pea family and native to humid lowland forested areas of South Asia. It features leathery medium-to dark-green compound leaves and is typically about 30-40 feet tall on average. The leaves are round to oval-shaped and drop in early winter, trifoliate (up to 10-18 inches wide), each having three rhombus-shaped leaflets held by a long petiole **(Figure 2.1 A)**. Black flower buds are formed in mid-winter on leafless stems; bicolor orange/red flowers are found (each about 2 inches long), blooming in dense clusters (racemes to about 6 inches long) from late January to March **(Figure 2.1 B)**. Flowers give way to fruits (flat, single-seeded pods up to 3-4 inches long), which emerge pale green but mature to bronze-brown.

**Figure 2.1.**
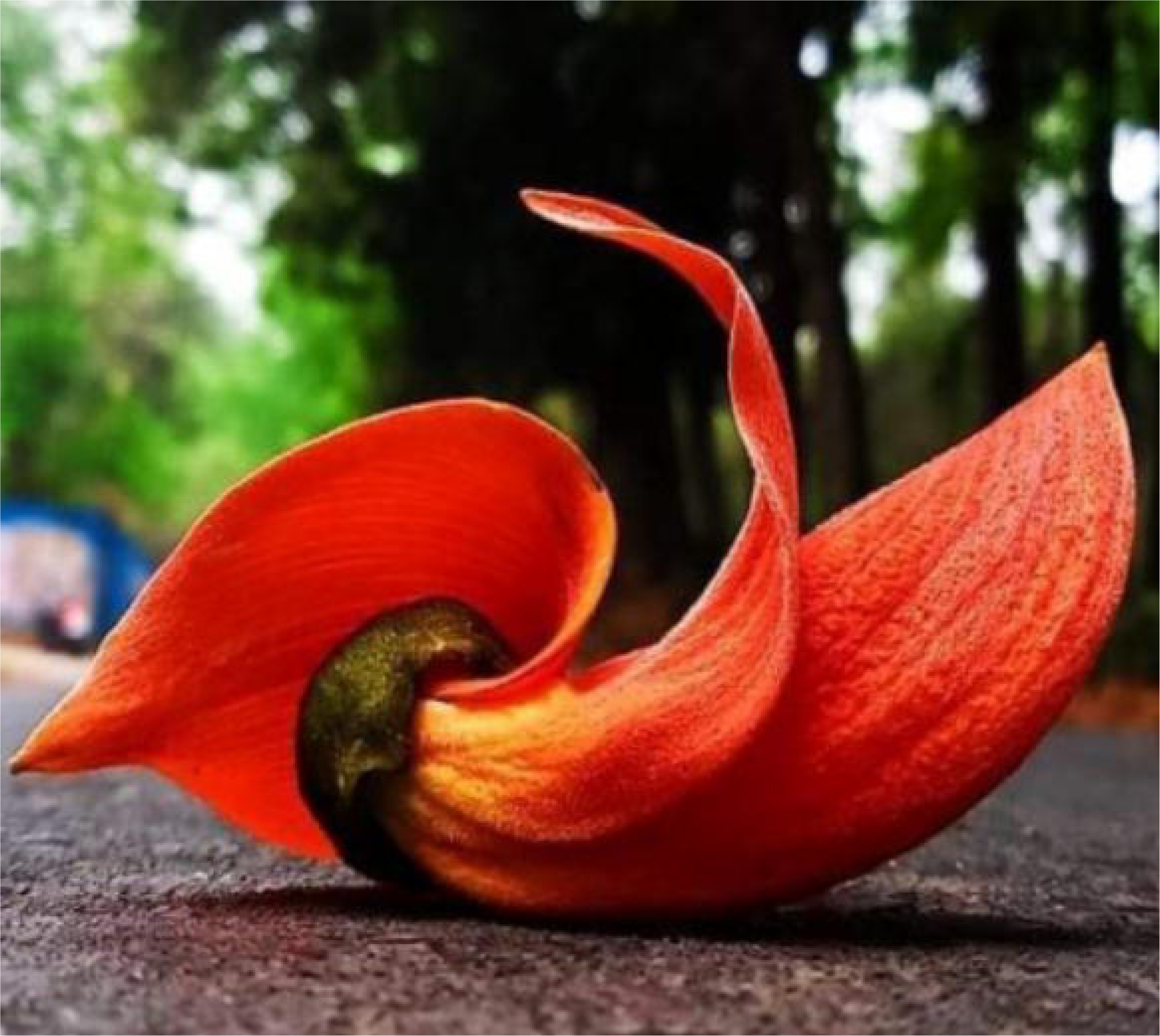
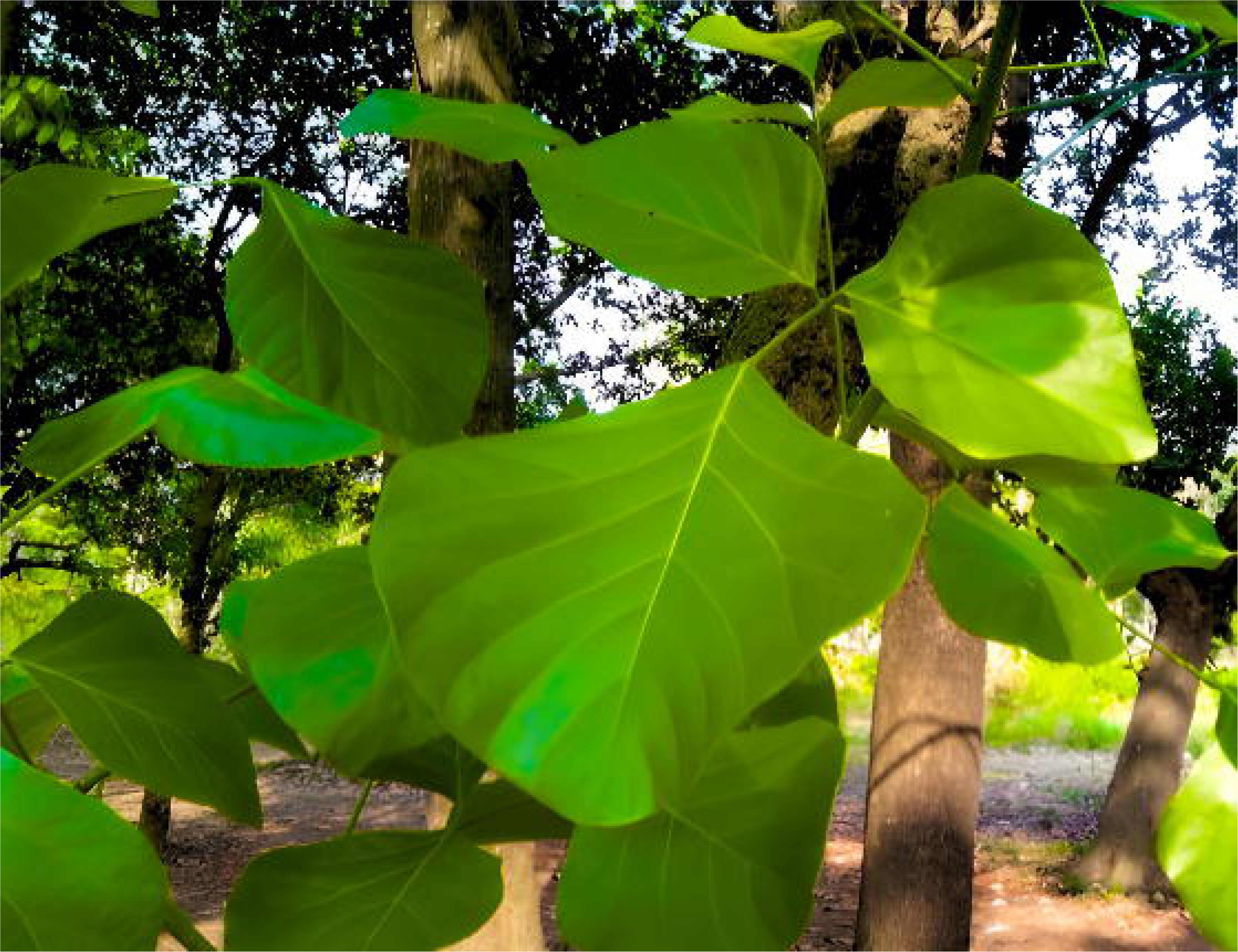
The morphology of *B. monosperma* (Flame of the Forest). **(A)** A close-up view of a distinctive orange-red flower of *B. monosperma.* **(B)** The characteristic large, trifoliate leaves and rough bark of the *B. monosperma* tree.

### 2. Collection and Preparation of Plant Material

#### Plant Collection

Plant parts (flower, bark, and leaf) of a locally grown, wild *B. monosperma* plant were collected from the Jahangirnagar University area, Savar, Dhaka, Bangladesh.

#### Drying and Pulverization

The collected raw plant materials were thoroughly washed with water to remove debris and other impurities. Subsequently, the different parts were sun-dried separately for 7 days until they reached a constant weight. The dried samples were then ground into a fine powder using an industrial grinder, followed by sieving to ensure uniform particle size. The resultant fine powders were weighed and stored separately before extraction.

### 3. Methanolic Extraction

One hundred grams (100g) of each powdered plant material (flower, bark, or leaf) were separately subjected to maceration in a volumetric flask containing 500 mL of 96% methanol as the solvent. The flasks were placed on a shaker and allowed to macerate for 72 hours at ambient room temperature and pressure to ensure complete extraction of the phytochemicals. Following the maceration period, the mixtures were filtered using Whatman Grade 1 filter paper (with a particle retention efficiency of 98% for 11 μm particles).

The filtered methanolic solutions were then concentrated by evaporating the solvent using a rotary evaporator (or equivalent apparatus) at a controlled temperature of 45°C under reduced pressure. This process was continued until the solvent was completely dried out, yielding the crude methanol extracts. The collected crude extracts were weighed to determine the extraction yield and stored in airtight containers at 4°C until required for further analysis.

### 4. Sample preparation for LC-MS/MS

An attempt to prepare a 1 ppm working solution for LC-MS/MS analysis was performed. Initially, a 500 ppm stock solution was prepared by accurately dissolving 5 mg of each individual crude extract (flower, bark, and leaf of *B. monosperma*) in 10 mL of 96% methanol (Sultana S and Asif HM, 2017). Subsequently, the 1 ppm working solution required for the analysis was created by performing a 1:500 dilution: a precise 0.02 mL aliquot of the 500 ppm stock was taken and diluted to a final volume of 10 mL with methanol. Afterward, the 1 ppm sample solution was sonicated (Powersonic 405, Hwashin technology Co., Korea) for 10 minutes in an ultrasonic bath to get the homogeneous mixture. This dilution step ( 10mL / 500ppm × 0.02mL = 1ppm) is critical for preventing column saturation and ensuring optimal ionization sensitivity during the LC-MS/MS run. A quaternary liquid chromatographic system (Nexara X2 series) with a triple quadruple mass spectrometer-mass spectrometer (LCMS-8050, Shimadzu Corporation, Japan) was used for sample analysis. The chromatographic separation utilized a mobile phase consisting of deionized water with 0.1% formic acid (Pump A) and acetonitrile with 0.1% formic acid (Pump B). The MS conditions were as follows: Run time: 2 minutes; Ion polarity: positive ion mode; ion source: electrospray ionization; Capillary voltage (kV): 5.0; Block temperature: 400°C; Desolvation line temperature: 200°C; CID gas: Argon (270 kPa), Nebulizing gas flow: N2, 1.5 L/min; Drying gas flow: N2, 10.0 L/min; Heating gas flow: 10 L/min; Interface temperature: 300°C; mass spectrum scan range (m/z): 200-400 Da. Previously prepared 1ppm samples of flower, bark, and leaf of B.monosperma were then loaded into the LC-MS chamber and set for a run.

### 5. *In-vivo* study

For the *in vivo* study, Swiss Albino-Wistar rats of both sexes were used as experimental animals. The rats were bred and maintained in the Animal Research Facility of the Department of Pharmacy, Jahangirnagar University. All experimental procedures were approved by the Ethical Committee of the Faculty of Biological Sciences, Jahangirnagar University, Savar, Dhaka, and conducted in accordance with internationally recognized ethical guidelines (NIH Publication No. 85-23, revised 1985). The animals were randomly divided into eleven groups, each consisting of three rats. Group I served as the control, Group II as the standard, and the remaining groups received three different doses (100 mg/kg, 200 mg/kg, and 400 mg/kg body weight) of extracts from three different plant parts—flower, bark, and leaf—administered via intraperitoneal injection.

#### Dose Preparation and Experimental Design

The plant extracts were diluted with distilled water (diH□O) to achieve the desired concentrations. Three different doses (100 mg/kg, 200 mg/kg, and 400 mg/kg body weight) were prepared for intraperitoneal administration. Before the experiment, all animals were fasted for 18 hours but had free access to water. Group I, or the control group, received normal saline orally at a dose of 25 ml/kg body weight. Group II was administered the same volume of normal saline containing Amlodipine + Olmesartan (5/20 mg) at a dose equivalent to 20 mg/kg body weight, serving as the standard. The combination of Amlodipine and Olmesartan (5/20 mg) is a well-established and effective antihypertensive therapy, represents an appropriate and ethically justified baseline antihypertensive therapy in studies evaluating the added effects of other agents, such as diuretics (Niemeijer & Cleophas, 2009). Groups III, IV, and V received the flower extract at doses of 100 mg/kg, 200 mg/kg, and 400 mg/kg body weight, respectively, dissolved in normal saline (25 ml/kg). Similarly, Groups VI, VII, and VIII received the bark extract, while Groups IX, X, and XI received the leaf extract at corresponding doses. All treatments were administered in such a manner that the total fluid volume per kilogram of body weight was consistent across all groups.

#### Urine Collection and Biochemical Analysis

Following dose administration, each rat was placed in an individual observation chamber and monitored for six hours at room temperature (25 ± 0.5°C). Urine samples were collected 30 minutes after dose administration and stored in glass collection tubes. The total urine volume was measured for each group, and concentrations of urinary electrolytes (Na□, K□, and Cl□) were determined using standard analytical methods. For Na□ and K□ estimation, 0.5 mL of freshly collected urine was digested with 8 mL of 69% HNO□ at 200 °C and 1600 atm, then diluted to 50 mL with distilled water. The concentrations were measured using a Shimadzu AA-7000 Atomic Absorption Spectrophotometer following AWWA 3111B (Na□) and AOAC 985.35 (K□) guidelines (Kazi et al.,2017; Poitevin, 2016). For Cl□ analysis, the Mercuric Thiocyanate Method was employed. One milliliter of urine was mixed with 100 µL of 0.25 M Ferric Ammonium Sulfate in 9 M HNO□ and 100 µL of saturated Mercuric Thiocyanate in ethanol. After 10 minutes of reaction, absorbance was measured at 460 nm using a UV-Vis spectrophotometer. Chloride concentration was determined from a calibration curve prepared with standard NaCl solutions (0–50 µg Cl□/mL).

## 3. Results

### 1. Analysis of LC-MS/MS

LC-MS/MS analysis of different parts of *B. monosperma* (flower,bark and leaf) revealed the presence of several clinically relevant diuretic compounds in the *B.monosperma* plant extracts. chlorthalidone, a thiazide-like diuretic, was consistently detected in all three plant parts (flower, leaf, and bark). In the positive ion mode of LC, chlorthalidone was identified by its diagnostic mass-to-charge (m/z) transitions of 339/135 and 339/195 **(Figure 4.1 A, B, C)**, which align with previously reported fragmentation patterns (Mazzarino et al., 2008). The widespread detection of chlorthalidone suggests that the plant naturally contains metabolites structurally similar to thiazide diuretics.

**Figure 4.1:**
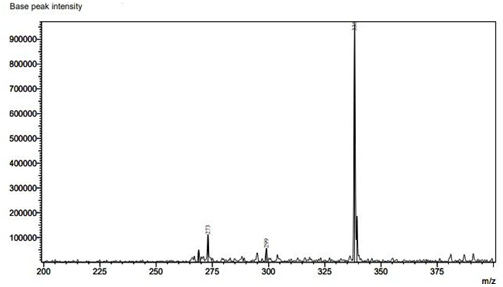
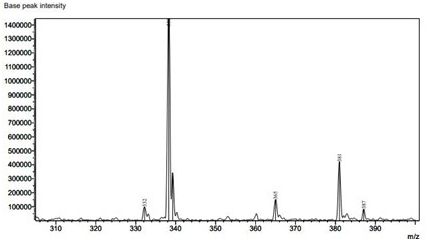
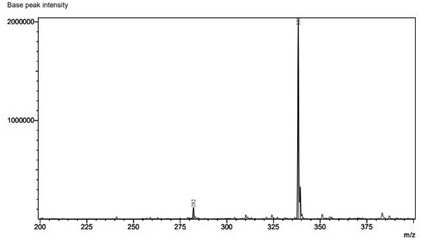
LC-MS/MS chromatogram of crude methanolic extract of different parts of *B. monosperma.* **(A)** Chromatogram of *B. monosperma* flower shows several distinct ion peaks, with notable signals detected around m/z ∼273, m/z ∼299, and a dominant major peak at approximately m/z ∼339, indicating the presence of high-abundance phytochemical constituents. Additional minor ions appear across the m/z 200–380 range, reflecting the complex metabolite profile of the flower extract. **(B)** The bark extract exhibits a highly intense major ion peak at m/z ∼336, accompanied by several medium-intensity peaks around m/z ∼332, m/z ∼365, m/z ∼381, and m/z ∼387, representing key secondary metabolites enriched in the bark. **(C)** The leaf extract exhibits a highly intensive major ion peak at m/z ∼ 336 and a medium intensive ion peak at m/z ∼ 282.

In addition, a trace amount of bumetanide was detected in the leaf and bark extracts, characterized by the transitions 365/184 and 365/240, consistent with bumetanide detection evident in **Figure 4.2 (B and C)** (G.P. et al., 2024). Interestingly, oxo-etozolin, a metabolite of etozolin, was observed exclusively in the flower extract, with transitions 299/202 and 299/127 confirming its identity **(Figure 4.2 A,B)**. Detection of oxo-etozolin is noteworthy, as it supports the potential of B.monosperma to produce or accumulate metabolites structurally related to synthetic diuretics (Mazzarino et al., 2008). Collectively, the results indicate that B.monosperma harbors bioactive metabolites structurally analogous to thiazide and loop diuretics, reinforcing its probable potential role in fluid balance regulation and validating its ethnopharmacological relevance.

**Figure 4.2:**
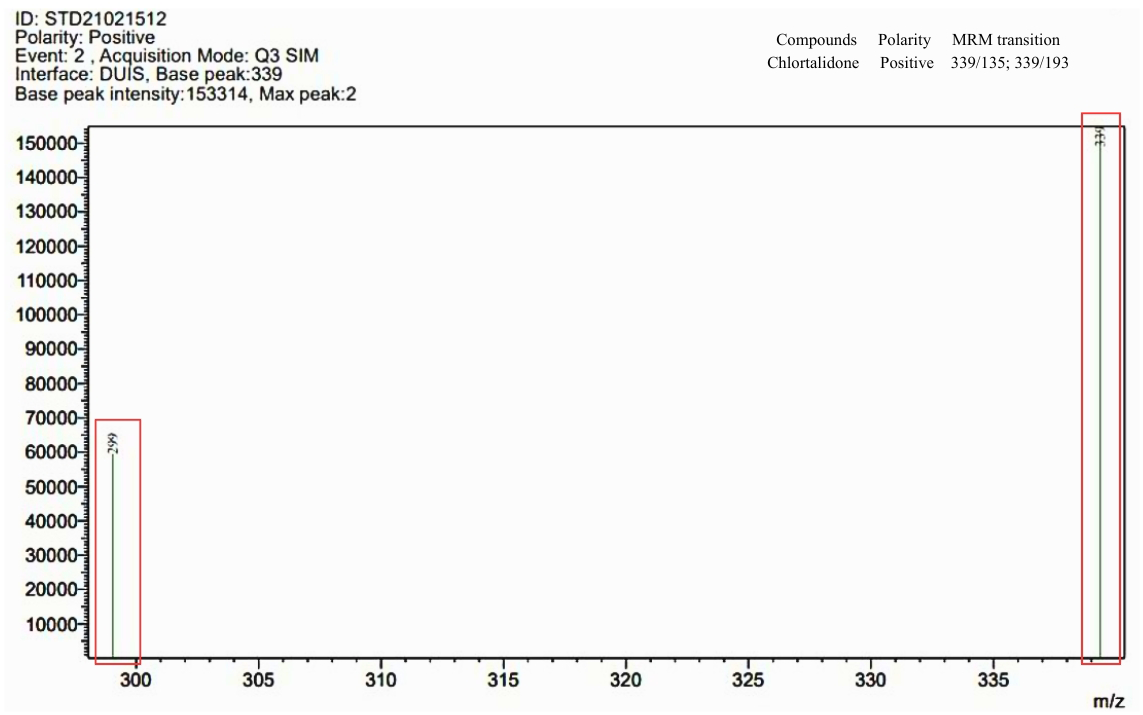
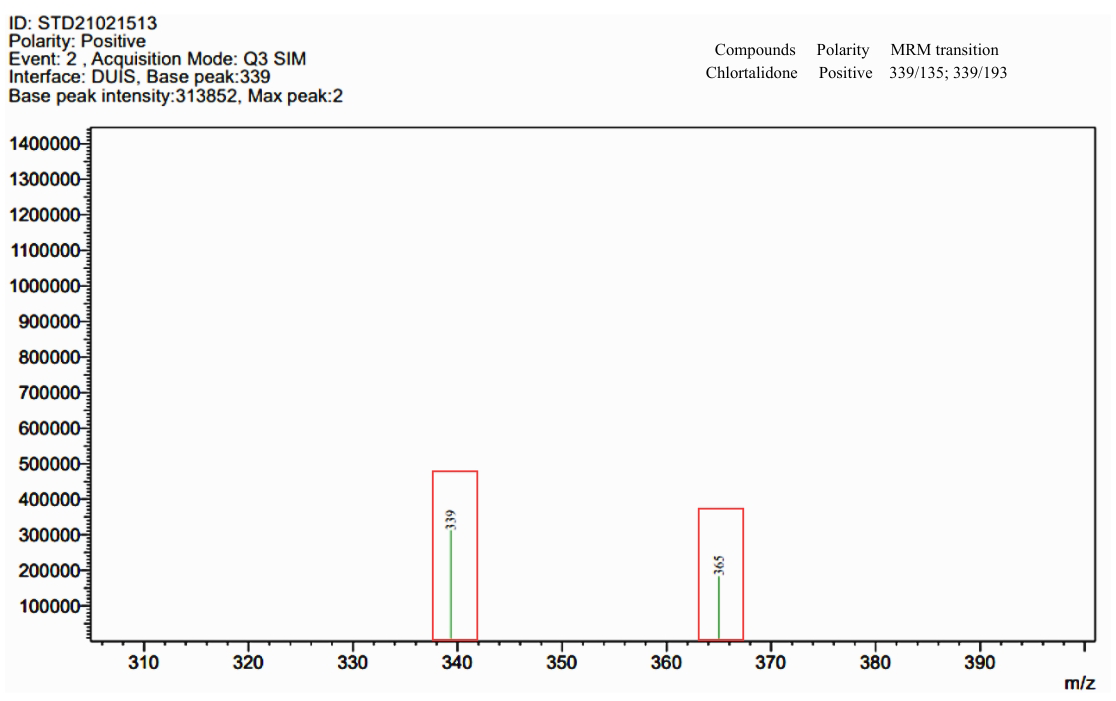
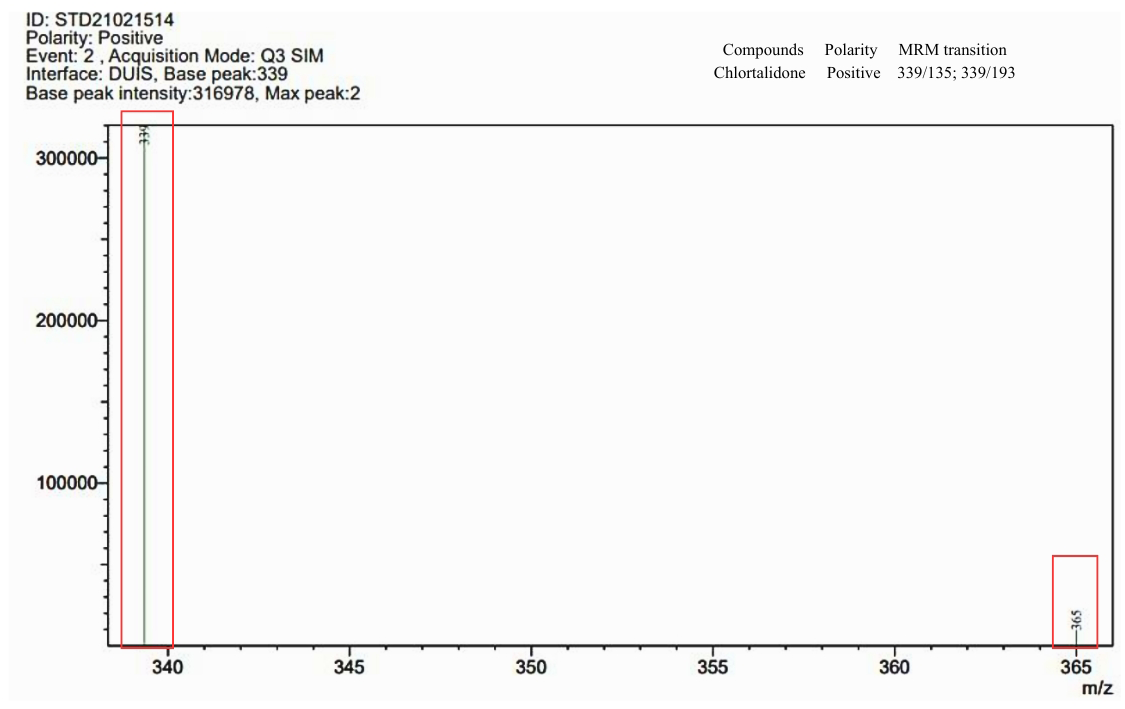
LC-MS/MS Chromatogram on Selective Ionization Mood of Different parts of *B. Monosperma.* **(A)** Flower where base peak 339 refers to Chlorthalidone and its intensity is 153314. **(B)** Bark where base peak 339 refers to Chlorthalidone and its intensity is 313852. **(C)** Leaf where base peak 339 refers to Chlorthalidone and its intensity is 316978.

#### Impact of Different Doses of *B. monosperma* Plant Parts on Urinary Electrolytes (Na□, K□, and Cl□), and Volume

The urinary electrolyte concentrations (Na□, K□, and Cl□) and total urine volume were measured to assess the diuretic potential of the plant extracts compared with the standard antihypertensive drug (Amlodipine + Olmesartan (5/20) mg) and control group. The results (**Table 4.1, Figure 4.3 A**) evidently revealed that administration of the standard drug (Amlodipine + Olmesartan (5/20 mg)) significantly increased urine output relative to the control group, indicating effective diuretic action.

**Table 4.1:** The collective urine volume of each experimental group after dose administration with accumulated data on concentration of Na^+^, K^+^ in urine samples for the test groups by AAS analysis and Cl^-^ from UV-vis spectrophotometry (unit in ppm).

| Name of Groups | Rat urine volume (mL) | Concentration of Na<sup>+</sup> (in ppm) in rat urine | Concentration of K<sup>+</sup> (in ppm) in rat urine | Concentration of Cl<sup>-</sup> (in ppm) in rat urine |
| --- | --- | --- | --- | --- |
| Control | 2.267±0.153 | 0.1688±0.0106 | 0.2414±0.0056 | 0.265±0.004 |
| Standard | 4.167±0.651 | 0.2238±0.0015 | 0.3704±0.001 | 0.767±0.036 |
| Group I (flower 100mg/Kg) | 2.767±0.208 | 0.2251±0.0065 | 0.2733±0.0123 | 1.018±0.005 |
| Group II (flower 200mg/Kg) | 3.567±0.513 | 0.3818±0.0085 | 0.1806±0.0055 | 0.892±0.006 |
| Group III (flower 400mg/Kg) | 3.6±0.4 | 0.4983±0.0127 | 0.2572±0.0078 | 0.961±0.005 |
| Group IV (bark 100mg/Kg) | 5.6±0.36 | 0.2384±0.0074 | 0.1459±0.0093 | 0.931±0.01 |
| Group V (bark 200mg/Kg) | 4.767±0.404 | 0.4938±0.007 | 0.1848±0.0098 | 0.990±0.006 |
| Group VI (bark<br>400mg/Kg) | 5.367±0.777 | 0.4081±0.0067 | 0.1228±0.0043 | 0.913.±0.007 |
| Group VII (leaf<br>100mg/Kg) | 2.037±0.503 | 0.9813±0.0141 | 0.3000±0.0106 | 0.899±0.007 |
| Group VIII (leaf<br>200mg/Kg) | 4.667±0.351 | 0.1775±0.0236 | 0.3785±0.0109 | 1.046±0.022 |
| Group IX (leaf<br>400mg/Kg) | 3.4±0.53 | 0.0347±0.0083 | 0.5155±0.0035 | 1.025±0.007 |

The results are expressed as mean±SD (n=3). Significant changes were found when all groups were compared with control and standard (p<0.05) using Tukey's Multiple Comparison Test in a one-way ANOVA

Similarly, other dose groups (Group I to IX) also exhibited an increase in urine volume, except for the leaf extract at 100 mg/kg (Group VII), although the degree of increase varied among the treatments. Among the plant parts tested, the bark extract demonstrated the highest diuretic activity, followed by the flower and leaf extracts. Within the flower extract groups, urine volume increased proportionally with dose, suggesting a dose-dependent diuretic response. In contrast, for the bark extract, the lowest dose (100 mg/kg) produced the greatest urine output, though the higher doses (200 and 400 mg/kg) also resulted in substantial diuresis. For the leaf extract, the 200 mg/kg dose group excreted the highest urine volume compared with the other leaf doses.

The urinary electrolyte concentrations (Na□, K□, and Cl□) and total urine volume were evaluated to determine the diuretic potential of the plant extracts in comparison with the standard antihypertensive drug (Amlodipine + Olmesartan, 5/20 mg) and the control group. As shown in **Table 4.1** and **Figure 4.3 A**, the standard drug produced a marked increase in urine output relative to the control, confirming its notable diuretic effect. Most extract-treated groups (Group I–IX) also demonstrated increased urine volume, with the exception of the leaf extract at 100 mg/kg (Group VII). Among the different plant parts, the bark extract exhibited the strongest diuretic activity, followed by the flower and leaf extracts. The flower extract showed a clear dose-dependent increase in urine output, whereas the bark extract displayed its maximum effect at the lowest dose (100 mg/kg), with higher doses still producing substantial diuresis. For the leaf extract, the 200 mg/kg group yielded the highest urine volume among the leaf-treated groups.

The accompanying graphs depict the urinary electrolyte concentrations (Na□, K□, and Cl□). For Na and Cl□, the control group showed comparatively lower levels than most treatment groups, except for Group IX (leaf extract, 400 mg/kg), as illustrated in **Table 4.1** and **Figures 4.3 B** and **4.3 D**. In contrast, urinary K□ levels were lower than the control in several groups, specifically Group II (flower extract, 200 mg/kg) and Groups IV, V, and VI (bark extracts at 100, 200, and 400 mg/kg), as presented in **Table 4.1** and **Figure 4.3 C**.

**Figure 4.3:**
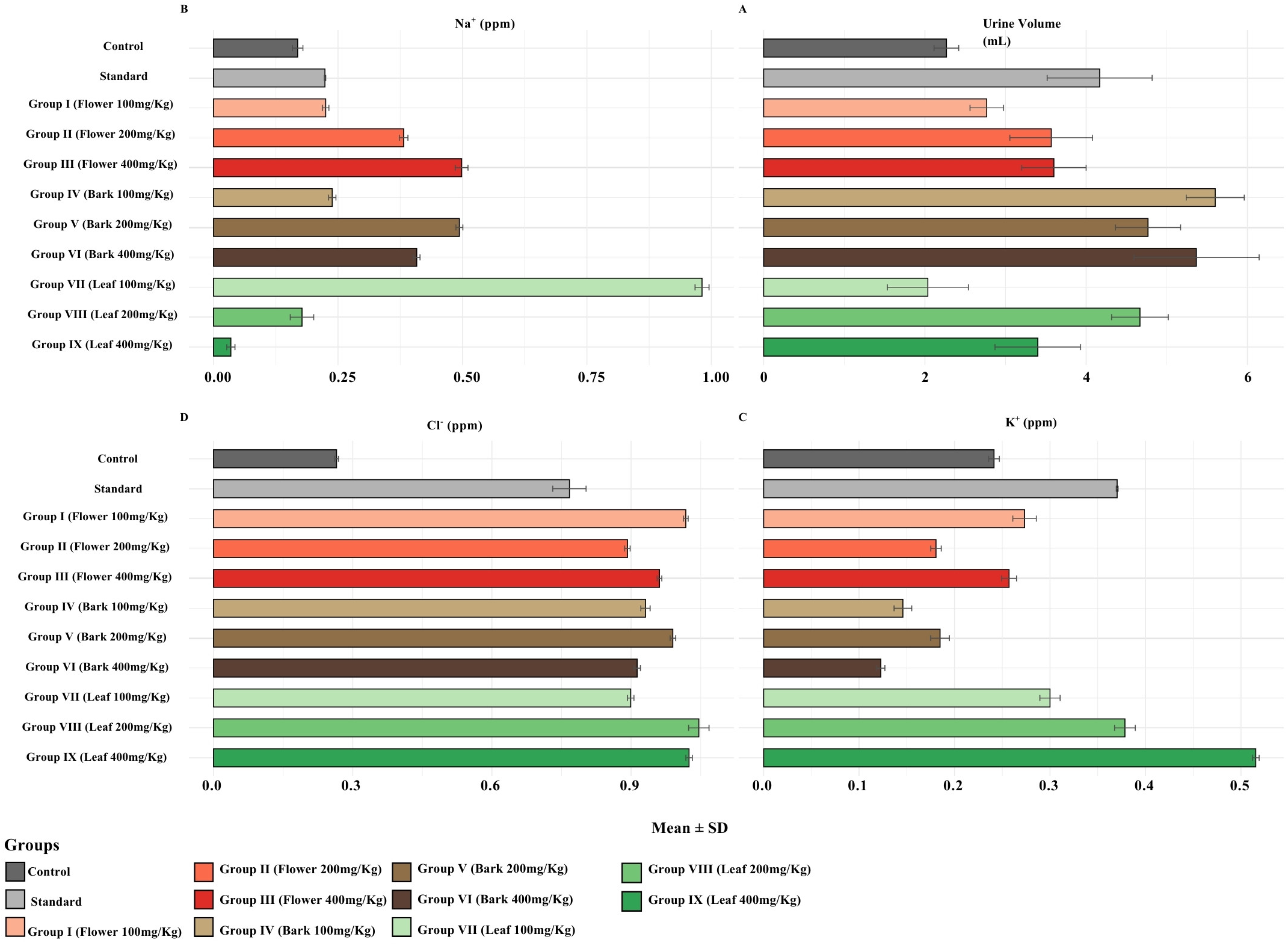
Effect of *B. monosperma* Extracts on Urine Volume and Electrolyte Excretion (Mean ± SD). **(A)** Urine volume (mL), **(B)** Sodium (Na□, ppm), **(C)** Potassium (K□, ppm), and **(D)** Chloride (Cl□, ppm) levels recorded in different experimental groups. Each horizontal bar represents the mean value for each parameter, with error bars indicating the standard deviation (SD). Treatments included methanolic extracts of *B. monosperma* flowers, bark, and leaves administered at 100, 200, and 400 mg/kg, compared with the Control and Standard groups.

#### Shift in Urine and Electrolyte Levels Compared to Control

Different parts of *B. monosperma* extracts represented marked differences in diuretic and saluretic responses among the plant parts **(Figure 4.4)**. The Standard drug (Amlodipine + Olmesartan (5/20) mg) produced a notable increased urine output (+1.900 mL) implying the diuretic effectivity in rats **(Figure 4.4 A)**. Among the extracts, the bark fractions imposed the strongest diuretic effect, with Group IV (100 mg/kg), Group V (200 mg/kg), and Group VI (400 mg/kg) showing increases of +3.333 mL, +2.500 mL, and +3.100 mL, respectively. Flower extracts generated moderate increases (+0.500 to +1.333 mL), whereas leaf extracts produced smaller or variable effects, ranging from a slight decrease in Group VII (–0.230 mL) to increases in Groups VIII and IX (+2.400 and +1.133 mL).

Changes in Na□ excretion **(Figure 4.4 B)** showed consistent increases in all Flower and Bark groups, ranging from +0.056 to +0.330 ppm. Bark extracts produced moderate elevations (+0.070 to +0.325 ppm). The highest Na□ excretion was recorded in Group VII (Leaf 200 mg/kg) at +0.81 ppm, followed by Group IX (+0.134 ppm). The Standard drug showed a mild rise (+0.055 ppm). The excretion of K□ **(Figure 4.4 C)** exhibited a distinct pattern, flower and bark extracts reduced K□, with values from –0.016 to –0.119 ppm, the lowest being Group VI (–0.119 ppm). Leaf extracts showed the opposite trend, with Group VII at –0.059 ppm, Group VIII at +0.137 ppm, and Group IX at +0.270 ppm. The Standard drug also increased K□ slightly (+0.129 ppm). Thus, only the higher-dose leaf extracts and the standard diuretic enhanced potassium elimination. All treatments increased chloride excretion **(Figure 4.4D)**. Flower extracts provided an increased excretion in the urination, between +0.627 and +0.753 ppm, while bark extracts showed similar elevations (+0.634 to +0.666 ppm). The highest Cl□ excretion occurred in the leaf groups, with Group VII (+0.781 ppm) and Group IX (+0.760 ppm) showing the strongest responses. The Standard drug produced a moderate increase (+0.502 ppm). Overall, the bark extract doses produced the most robust diuretic effect, while leaf extracts particularly at higher doses evidently showed the strongest saluretic (Na□ and Cl□) responses and a dose-dependent rise in K□ excretion. Flower extracts demonstrated moderate but consistent activity across all parameters.

**Figure 4.4:**
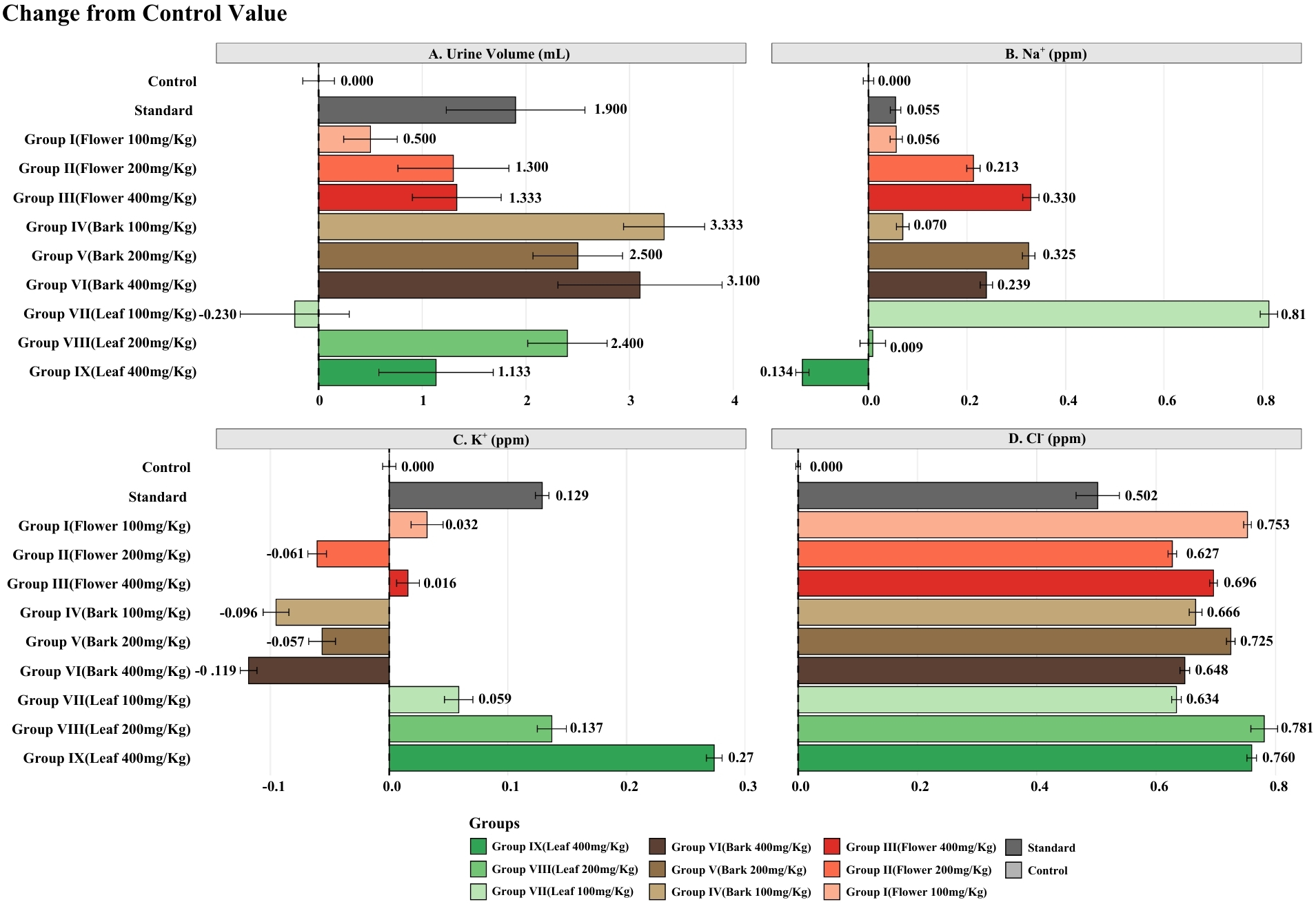
Shift from Control Value for Urine Volume and Electrolyte Excretion in Experimental Groups. The plot displays the Mean ± Standard Deviation (SD) of the change in **(A)** Urine Volume (mL), **(B)** Sodium (Na+) (ppm), **(C)** Potassium (K+) (ppm), and **(D)** Chloride (Cl−) (ppm) for the various treatment groups relative to the Control group (set at 0.000). Treatments include a Standard drug and nine different plant extract groups (Groups I–IX) administered at various concentrations (mg/Kg). Positive values indicate an increase, while negative values indicate a decrease in the measured parameter compared to the Control..

## 5. Discussion

The present study investigated the diuretic activity of different parts of *Butea monosperma* (flower, bark, and leaf) using an in vivo rat model, alongside LC-MS/MS analysis to identify potential bioactive compounds responsible for this pharmacological action. The findings revealed a dose-dependent increase in urine output and electrolyte excretion, with the bark extract demonstrating the most pronounced effect compared to the flower and leaf extracts. From the LC-MS/MS study, the findings revealed the presence of chlorthalidone, which has been shown to have great potential for anti-hypertensive efficacy found in several prospective studies, and poses a long half-life, especially in combination with multiple antihypertensive drugs (Ishani A et al., 2022). The question arises of how this helps in other ways, from plant extracts towards medicinal consumption.

The increase in urinary volume and electrolyte excretion (Na□, K□, and Cl□) observed later in treated rats implies that the plant extracts may influence renal tubular reabsorption mechanisms, similar to conventional diuretic agents. Among the extracts, the bark exhibited the strongest effect at 200 mg/kg and 400 mg/kg doses, showing comparable or even superior diuretic performance to the standard antihypertensive treatment. And earlier in the LC-MS/MS study and from analysis, chlorthalidone was identified. Chlorthalidone is a thiazide-like sulfonamide-derived diuretic and has been FDA-approved since 1960 to manage hypertension (P. Patel & J. B. Patel, 2024). It is a first-line agent for the treatment of hypertension, utilized both as an isolated agent and in combination with other antihypertensive drugs, including β-blockers or clonidine (V. Thanikgaivasan, 2017). It is a very commonly used diuretic found in nature, and works as an anti-hypertensive by retaining excess water along with electrolytes (i.e: Na^+^, K^+^, Cl^-^) from blood and excreting them through urination (Whelton et al., 2018). The increase in urinary volume and electrolyte excretion (Na□, K□, and Cl□) observed in treated rats implies that the plant extracts may influence renal tubular reabsorption mechanisms, similar to the conventional diuretic agents. Among the extracts, the bark exhibited the strongest effect at 200 mg/kg and 400 mg/kg doses, showing comparable or even superior diuretic performance to the standard antihypertensive treatment. Electrolyte analysis showed concurrent increases in Na□, K□, and Cl□ excretion, indicating that *B. monosperma* extracts act through a mechanism resembling that of loop diuretics or thiazide diuretics rather than potassium-sparing agents (Sundaresan et al., 2017). This pattern of electrolyte loss, while pharmacologically beneficial in managing hypertension and edema, also emphasizes the need for dosage optimization to prevent hypokalemia or excessive fluid loss during therapeutic use (Khow et al., 2014). Compared to synthetic diuretics, the extracts showed fewer adverse behavioral effects in the test animals, suggesting a safer pharmacological profile. This supports the potential of *B. monosperma* as a natural diuretic candidate for further development. However, while the bark extract displayed strong efficacy, variations in potency between plant parts highlight the importance of standardized extraction and compound quantification in future pharmacognostic research. In this study, it is evident that the different parts (flower, bark, and leaf) of the plant have intensified the containment of chlorthalidone. The leaf and bark part has a higher intensity for the base peak for chlorthalidone and can be concluded as the more efficient effects for the diuretic or anti-hypertensive effect upon consumption. Overall, the results of this investigation provide scientific evidence that *Butea monosperma*, particularly its bark extract, exhibits potent diuretic properties through dose-dependent stimulation of urine and electrolyte excretion

From ancient times, *B. monosperma* has been extensively used in the treatment of a wide array of diseases, most preferably in the Indian subcontinent (Singh & Srivastava, 2022). Worms, constipation, piles, diabetes, and a clogged throat have all been treated with *B. monosperma* in the past (Iqbal et al., 2006). A report by Bandara et al. (1989) recorded the use of bark for the treatment of diarrhea, dysentery, ulcers, sore throat, diabetes, polypus in nose and snake bite. Also, round worms are treated with B. monosperma pods (Rajbhandari et al., 2000). *B. monosperma*, as a medicinal plant have been long recognized and promising, found to be a source of bioactive molecules for developing novel agents for the prevention and treatment of various diseases (Subramaniyan B., Kumar V., & Mathan G., 2017). Within the scope of knowledge, plant remedies have been more beneficial and less harmful than synthetic ones, proven in many cases. Under adverse environmental conditions, plants have the ability and need to generate stress by synthesizing species-specific secondary metabolites under sudden changes in rain, drought, humidity, temperature, and infections by phytopathogens (Erb M & Kliebenstein DJ, 2020). Likewise, *B. monosperma* has high tolerance to adverse environmental conditions by maintaining its physiological parameters, indicating alterations in metabolic functions, enzyme activities, antioxidant productions, and membrane damage due to pollution stress. The major secondary metabolites are flavonoids, such as butyrin, isobutyrin, butein, isocoreopsin, and chalcone, which impart valuable medicinal properties to this plant (Hiremath K. Y., Veeranagoudar D. K., & Bojja K. S., 2024). But metabolites have demonstrated antiproliferative activity, immunomodulatory effects, and modulation of cell signaling; their full mechanism of action is yet to be evaluated. Likewise, from this study, our deduction in this perception that the evidence of diuretics, thiazide-like compound chlorthalidone, provides a head start on whether it is efficient to use the tree parts of *B. monosperma* directly for hypertension treatment. This experiment not only led to the existence of the treatment, but the efficacy was clinically challenging. Several studies included an *in vivo* study on albino rats, which was performed to literally understand its medicinal values in other perspectives. One study showed the toxic effect of seed powder from *B. monosperma* when administered in powder form; though this study used dosage was not found to be so extreme (Behera B., Pradhan S., Samantaray A., & Pradhan D., 2020).

Finally, our study findings support the traditional medicinal claims and suggest that the bark extract could be explored as a natural alternative or adjunct to conventional diuretics for hypertension and related cardiovascular conditions. Future studies should focus on isolating and characterizing the specific bioactive molecules responsible for the diuretic effect, elucidating their mechanisms of action, and assessing long-term safety and efficacy through detailed pharmacological and toxicological evaluations.

## Conclusion

*B. monosperma* possesses an efficient indigenous origin and remarkable research potential, requiring consistent, validated documentation and further definitive studies to explore its pharmacological promise. Our findings suggest that it contains diuretic compounds, specifically thiazide-like agents, that probably contribute to a higher chances to its natural antihypertensive properties. The bark extract exhibited the most potent effect, producing the highest urine volume and electrolyte (Na□, K□, Cl□) excretion, closely comparable to the standard antihypertensive treatment. These findings scientifically validate the traditional use of *B. monosperma* in the management of hypertension and fluid retention. The observed activity is likely attributed to the presence of bioactive phytochemicals such as flavonoids, alkaloids, and saponins, which enhance renal excretion and fluid balance. Overall, *B. monosperma*, particularly its bark extract, shows promising research focusing on the isolation of active compounds, elucidation of mechanisms of action, and detailed safety profiling. It is recommended to support its future development as a phytopharmaceutical candidate.

## Supporting information

Mice feeding and weight data

## Acknowledgement

All the chemical tests were done at Bangladesh Reference Institute for Chemical Measurements (BRiCM), Dr. Kudrat-i-Khuda Road, Dhanmondi, Dhaka-1205, Bangladesh. A special thanks to Md. Moniruzzaman (Senior Scientific Officer, BRiCM) for his special support and encouragement. Also, to Pranab Karmakar, PhD (Senior Scientific Officer, BRiCM) Md. Abu Hasan (Scientific Officer, BRiCM), and Mahmudul Hasan Razu (Scientific Officer, BRiCM) for their kind consultancy. Also, a humble gratitude to Esha Binte Shahriar, Aminul Islam and Rubiat Afrin (Laboratory of Preventive and Integrative Biomedicine, Department of Biochemistry and Molecular Biology, Jahangirnagar University, Savar, Dhaka-1342, Bangladesh) for their support.

